# Evapotranspiration is resilient in the face of land cover and climate change in a humid temperate catchment

**DOI:** 10.1101/075598

**Authors:** S. K. Hamilton, M. Z. Hussain, C. Lowrie, B. Basso, G. P. Robertson

## Abstract

In temperate humid catchments, evapotranspiration returns more than half of the annual precipitation to the atmosphere, thereby determining the balance available to recharge groundwaters and support stream flow and lake levels. Changes in evapotranspiration rates and therefore catchment hydrology could be driven by changes in land use or climate. Here we examine the catchment water balance over the past 50 y for a catchment in southwest Michigan covered by cropland, grassland, forest, and wetlands. Over the study period about 27% of the catchment has been abandoned from row-crop agriculture to perennial vegetation and about 20% of the catchment has reverted to deciduous forest, and the climate has warmed by 1.14°C. Despite these changes in land use, precipitation and stream discharge, and by inference catchment-scale evapotranspiration, have been stable over the study period. The remarkably stable rates of evapotranspirative water loss from the catchment across a period of significant land cover change suggest that rainfed annual crops and perennial vegetation do not differ greatly in evapotranspiration rates, and this is supported by measurements of evapotranspiration from various vegetation types based on soil water monitoring in the same catchment. Compensating changes in the other meteorological drivers of evaporative water demand besides air temperature—wind speed, atmospheric humidity, and net radiation—are also possible, but cannot be evaluated due to insufficient local data across the 50-y period. Regardless of the explanation, this study shows that the water balance of this landscape has been resilient in the face of both land cover and climate change over the past 50 y.

## 1. Introduction

In temperate humid catchments, evapotranspiration (ET) returns more than half of the annual precipitation to the atmosphere (Hanson, 1991; Williams *et al.,* 2012; Zhang *et al.,* 2016), mainly during the growing season by plant transpiration. The balance between precipitation and ET recharges groundwaters and supports stream flow and lake levels. Paired catchment studies often have shown that changes in the nature of the vegetation cover, especially deforestation or afforestation, alter ecosystem ET rates and thereby change stream flows (Bosch and Hewlett, 1982; Hornbeck *et al.,* 1993; Zhang *et al.,* 2001; Price, 2011; Brown *et al.,* 2005 and 2013). However, these studies are often conducted in small experimental catchments and generally compare stream water yields between two kinds of perennial vegetation (woody and herbaceous).

There have been fewer catchment-scale comparisons of water yield from annual vegetation such as maize (*Zea mays*, known as corn in the U.S.) and soybean (*Glycine max*) vs. perennial vegetation such as forest or grasslands (Price, 2011), yet land cover change from perennial vegetation to cropland and vice versa has occurred throughout the world as a result of agricultural expansion and contraction. In eastern North America, the original forests and grasslands were largely converted to agricultural lands by European settlers, but since the mid 1900s a substantial fraction of the converted land has reverted back to successional fields and forests as the more marginal agricultural lands were abandoned due to low profitability, poor suitability to mechanized cultivation, and concerns about soil erosion and degradation (Houghton and Hackler 2000; Ramankutty *et al.,* 2010).

Land cover in agricultural regions is expected to continue to change in the future. As grain crops have become more profitable over the past decade due to global demand for food and US policies that support ethanol production from maize, more land in grasslands (including CRP land) is being converted to grow maize and soybean (Lark *et al.,* 2015). Meanwhile, successional ecosystems are becoming mature forests in many locations (Pugh, 2015). Climate change and invasive plant species will increasingly drive changes in the nature and phenology of vegetation communities (Simberloff, 2000; Parmesan and Hanley 2015). Further changes to the nature of vegetation in agricultural landscapes may occur if cellulosic biofuel crops are increasingly grown in the future (Gelfand *et al.,* 2013).

Recently we reported ET measurements in candidate cellulosic cropping systems at a location in southwest Michigan, USA using two distinct approaches: 1) by monitoring soil water content with time domain reflectometry in annual crops (maize) as well as perennial grasslands and hybrid poplar stands (Hamilton *et al.,* 2015); and 2) by monitoring energy and water vapor fluxes using eddy covariance in maize, switchgrass, and prairie at a nearby site (Abraha *et al.,* 2015). Results suggest strikingly similar growing-season ET among these diverse plant systems, raising the question of whether land cover changes would significantly affect ET in the Midwest U.S., as suggested in some modeling studies (e.g., Le *et al.,* 2011; VanLoocke *et al.,* 2012; Zhuang *et al.,* 2013).

The objective of this study is to examine trends in ET over 50 y in a particularly well-characterized, temperate humid catchment that has experienced significant land cover change, but without the complications of urbanization, dams, and stormwater management changes that are typical of larger catchments. We infer ET from the balance between precipitation and discharge, and the results are compared with our independent measurements of ET made on annual and perennial vegetation in the same catchment.

## 2. Methods

### 2.1 Study site

Augusta Creek is a 3^rd^-order stream in southwest Michigan (Kalamazoo and Barry counties) that drains a predominantly rural landscape (95 km^2^) composed of a mosaic of forest, fallow fields, annual crops, wetlands, lakes, light residential development, and golf courses (Figure 1). There are no impervious surfaces or storm drainage systems that drain into the stream above the discharge measurement point, and urban land use covers just 2.4% of the catchment (land cover proportions over the 50-y period are presented later). The stream is groundwater-fed, gaining water along most of its length. Its tributaries emanate from wetlands or small lakes, and prairie fen wetlands line much of the stream channels.

**Figure 1.**
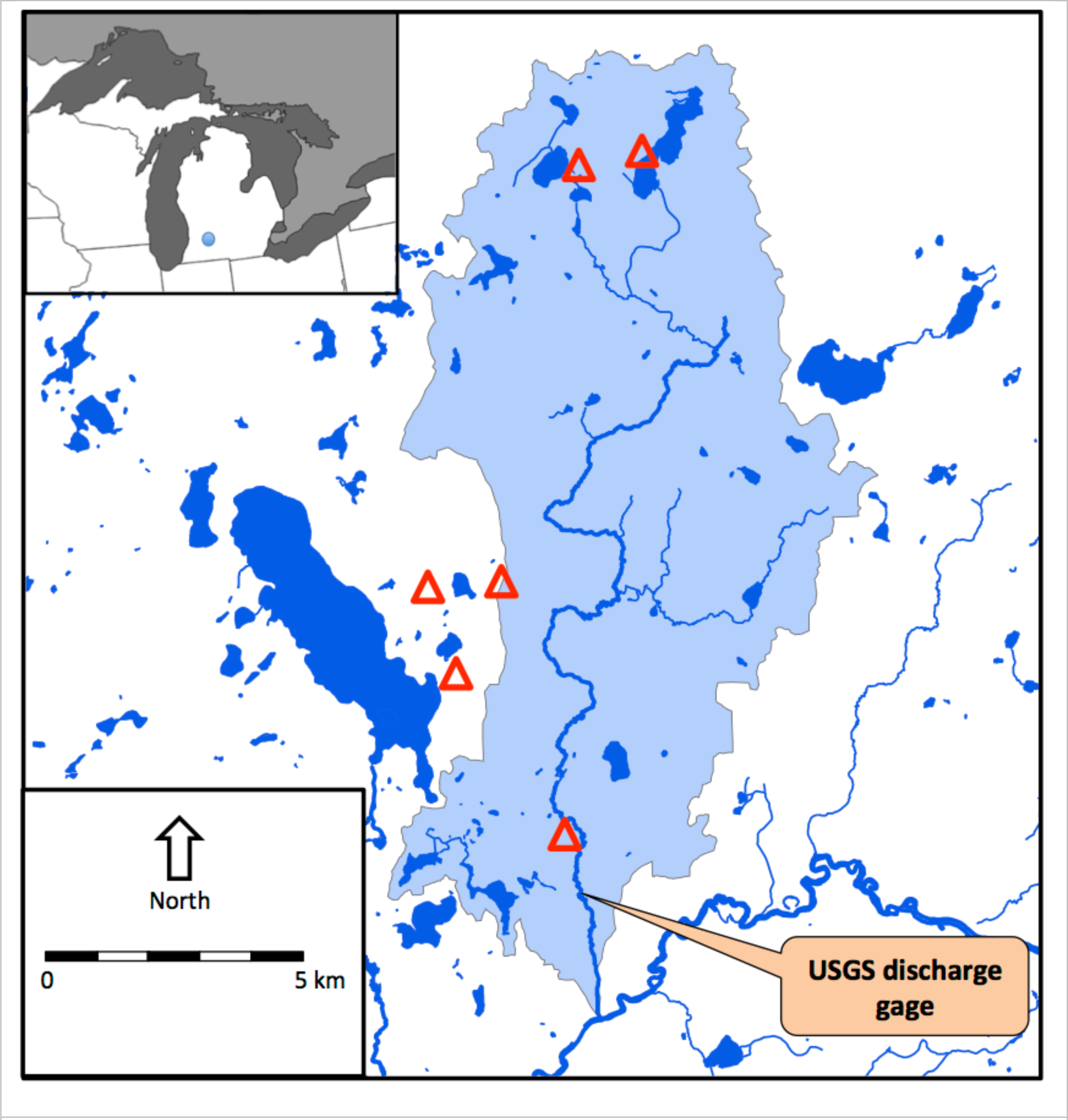
Location of Augusta Creek in Michigan (inset) and watershed boundaries (shaded). Precipitation measurement sites are shown by triangles.

The stream runs through deep glacial deposits that lie well above the bedrock. The most common soils in upland areas are well-drained Typic Hapludalfs developed on postglacial terrain (Thoen, 1990), and there is little to no overland flow from upland areas to the stream due to the high permeability of these coarse-textured soils (Rheaume, 1990). Irrigation of crops was rare in the area until very recently; some expansion has taken place since 2005, supplied by groundwater wells.

Augusta Creek is in the vicinity of the W.K. Kellogg Biological Station (KBS), where we conduct agricultural experiments under the aegis of the Great Lakes Bioenergy Research Center (GLBRC) and KBS Long Term Ecological Research site (www.lter.kbs.msu.edu; 42.3956° N, 85.3749° W and 288 m asl). Mean annual air temperature is 10.1 °C and annual precipitation is 1005 mm, 511 mm of which falls as rain during the May-Sep growing season (1981–2010) (NCDC 2013). In this region ET is normally water–limited during the warmer part of the year (i.e., during at least part of the growing season), and energy–limited during the cooler months (McVicar *et al.*, 2012a).

### 2.2 Land cover changes

Land cover for 1960 was estimated from georectified and mosaicked aerial photographs in a geographic information system (ArcGIS). The catchment boundaries above the discharge measurement point (US Geological Survey; Hydrologic Unit Codes 04050003040060 plus 04050003040070) and wetlands and lakes (National Wetlands Inventory: http://www.fws.gov/wetlands/) were overlain on the aerial photo mosaic and land cover was examined in the upland portions of the catchment. Based on the National Wetlands Inventory, wetlands and lakes contiguous with the stream system amount to 15.9 km^2^, or 16.6 % of the catchment above the discharge measurement point. Isolated wetlands and small lakes also occur throughout the upland catchment, covering 5.2% of its area. Wetland areas were assumed to be constant over the study period; there has been no wetland drainage or creation in the catchment since 1960, and within the area mapped, wetland boundaries generally include intermittently wet soils with high water tables as well as areas with surface water.

Land cover for 2014 was estimated from the Cropland Data Layer (http://nassgeodata.gmu.edu/CropScape/). For this purpose, we combined all field crops (primarily maize, soybean and small grains) into the annual crop category, and all forests (deciduous and coniferous) into the forest category. Conifers are not native to the upland landscape here, but have been planted throughout the catchment; their total area as of 2014 amounts to ∼3% of the total forest area and 1% of the upland catchment. The grassland and pasture category includes hay as well as fallow fields (no native grassland remains). The width of rural roads was exaggerated 3–5 fold in the Cropland Data Layer, presumably due to automated classification of mixed pixels, so vegetated edges of roadways were manually reclassified as grasslands. Land cover for an intermediate date (1978), based on aerial photo interpretation, was available from the Michigan Resource Inventory System (MIRIS) (http://www.ciesin.org/IC/mdnr/mrip.html); this data set was comparable for forest but combines annual crops with some kinds of pasture, and was therefore not compared for those categories.

### 2.3 Discharge and climate records

The US Geological Survey has monitored the discharge of Augusta Creek below the lowermost tributary inflow since 1964 (station 04105700). The long-term mean discharge at this point, which drains 95.3 km^2^, is 1.28 m^3^ s^-1^. Daily discharge measurements for October 1964 through September 2014 were partitioned into baseflow and stormflow using the Web-based Hydrograph Analysis Tool (WHAT) described by Lim *et al.*, (2005). Mean annual baseflow and stormflow discharges were calculated on a standard U.S. water-year basis beginning on 1 October of each year, representing the transition between warm and cool seasons, and water years are labeled by the starting year (i.e., water year 1964 is 1 October 1964–30 September 1965).

Climate data were drawn from several sources and compiled on a water-year basis. Precipitation observations are from at least three stations (except 1992 which has two) distributed across the catchment from north to south (Figure 1). Air temperature, saturated vapor pressure, and drought index data were obtained from the Midwest Regional Climate Center (http://mrcc.isws.illinois.edu/).

### 2.4 Estimation of evapotranspiration from water balances

Evapotranspiration has often been estimated from catchment water balances (e.g., Zhang *et al.,* 2016). For Augusta Creek, the water balance for the upland portion of the catchment was determined as the difference between annual totals of precipitation falling on the uplands (i.e., the catchment excluding wetlands and lakes contiguous with the stream channels) and the annual stream baseflow discharge. Isolated lakes and wetlands were included in the upland catchment area. The difference between precipitation inputs on the uplands and stream baseflow outputs is therefore considered to represent the ET of the upland catchment.

This approach to ET estimation assumes that stormflow represents direct capture of precipitation from the wetlands and lakes contiguous with the stream system, whereas baseflow represents infiltration and percolation of precipitation falling on the upland catchment. The validity of this assumption is supported by the water balance calculations (see Results below) as well as the high permeability of the soils in the uplands (Rheaume, 1990). Other assumptions that are reasonable in this case include no inter-basin transfers of water, which is true in this catchment, and no significant trend in water storage in the aquifer or surface water bodies over the study period. Although there are no continuous water table measurements spanning this study period, water levels of local lakes that are connected to the groundwater have shown no unidirectional trend since the late 1960s (see Figure S1 for an example of water level data for a lake in the Augusta Creek catchment). Additional evidence for no interannual trend in groundwater levels is provided by a compilation of static water level measurements that are made when residential water supply wells are constructed, which shows no trend over the study period (Figure S2).

### 2.5 Evapotranspiration estimation from soil water content measurements

Since 2009, soil water profiles throughout the root zone and below were monitored hourly using permanently installed, horizontally inserted time domain reflectometry (TDR) probes at depths of 20, 35, 50, 65, 90 and 125 cm as well as a vertically inserted probe at 0–10 cm depth. Our methods for estimating ET from soil water profiles are described by Hamilton *et al.* (2015), who presented data on six biofuel cropping systems harvested each fall. The TDR measurements provide an estimate of ET when daily drawdowns in soil water can be measured and the soil water content is below its drained upper limit, which is typical of most of the growing season. The sum of the daily drawdowns in soil water content over the entire profile (0-150 cm) across the growing season provides an estimate of ET; on days when new infiltration of rain water prevented a measurable soil water drawdown, we estimated ET using a crop growth model (Basso and Ritchie, 2012).

Here we present the mean ET rates for three of those systems that resemble vegetation found on the broader landscape: 1) continuous no-till maize; 2) a restored native prairie planted with 18 species of forbs and grasses; and 3) a hybrid poplar plantation (*Populus nigra × P. maximowiczii ‘NM6’*). In addition, we present comparable water use measurements for three other systems in the same vicinity: 1) a fallow field abandoned from row-crop agriculture in 2008 and harvested each fall; 2) a mature deciduous forest (>50 y old) dominated by sugar maple (*Acer saccharum*), red oak (*Quercus rubra*) and hickory (*Carya* spp.) trees; and 3) an early successional forest (ca. 25 years old) dominated by shrubs including autumn olive (*Elaeagnus umbellata*) and honeysuckle (*Lonicera* sp.) as well as a few medium-sized sugar maple and black cherry (*Prunus serotina*) trees.

## 3. Results

### 3.1 Land use and climate changes

Maize has been the dominant agricultural crop over the 50-y study period with the balance of harvested crops shifting increasingly to soybean since the 1970s, as in the greater Midwest US region (Gage *et al.,* 2015). Data on Kalamazoo County from the annual Census of Agriculture (U.S. Department of Agriculture: http://www.agcensus.usda.gov/) indicate that in 1964 maize accounted for 69% of harvested cropland, soybean for 5.7%, and the balance was mostly oats with some barley and wheat. By 1987 maize was 58% and soybean 28% of harvested cropland, and by 2007 these two crops accounted for 64% and 32% of the harvested cropland.

Land cover in the upland catchment changed significantly between 1960 and 2014 (Figure 2). The proportion of the upland catchment in annual crops decreased from 57 to 30%, while forest increased from 15 to 35%. The proportion of grassland remained similar, although only 20% of the 1960 upland grassland was still grassland in 2014; most of the 1960 grassland became forest (43%) or cropland (22%), while some newly abandoned cropland became grassland. The 1978 MIRIS land cover data (not shown; see Methods) indicate that 94% of the forest present in 2014 existed by 1978, so most reforestation began between 1960–78. Urban and residential development represents a small fraction of the catchment (<2.4%), not including golf courses created during the study period that covered 4.5% of the upland catchment by 2014 (the golf courses occasionally irrigate during dry summers but are not significant water users at the catchment scale). Similar changes in land cover occurred in adjacent catchments.

**Figure 2.**
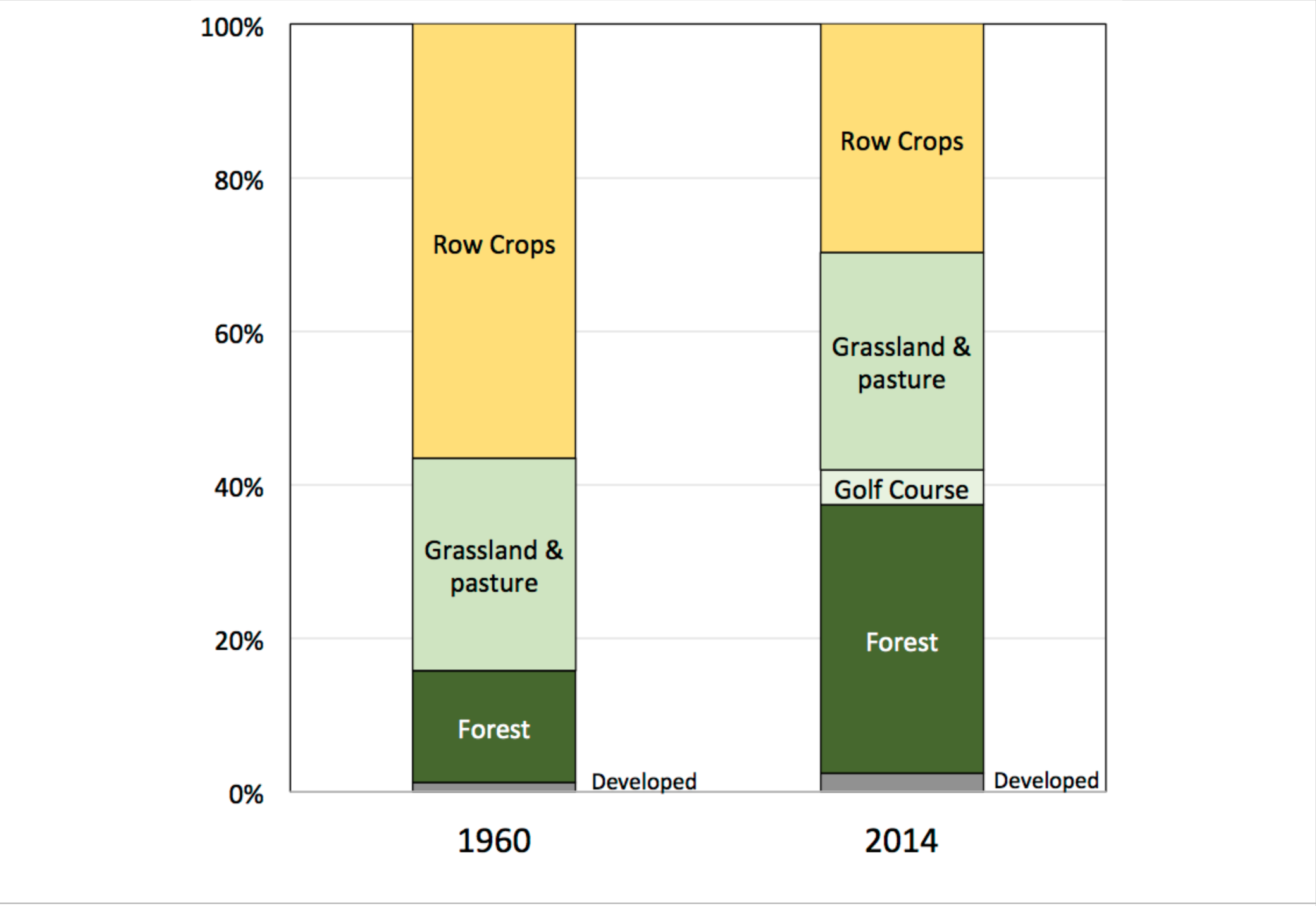
Land cover change in the Augusta Creek watershed. Estimates are based on analysis of aerial photos (1960) or satellite imagery (2014 Cropland Data Layer: http://nassgeodata.gmu.edu/CropScape/).

Annual precipitation for the Augusta Creek catchment over the 50 y averaged 948 ± 118 mm y^-1^ with no linear temporal trend (p = 0.93) (Figure 3a). No linear trend exists in mean annual values for either the Palmer Drought Severity Index or the Palmer Hydrological Drought Index (p = 0.34 and 0.67, respectively; Figure S3).

**Figure 3.**
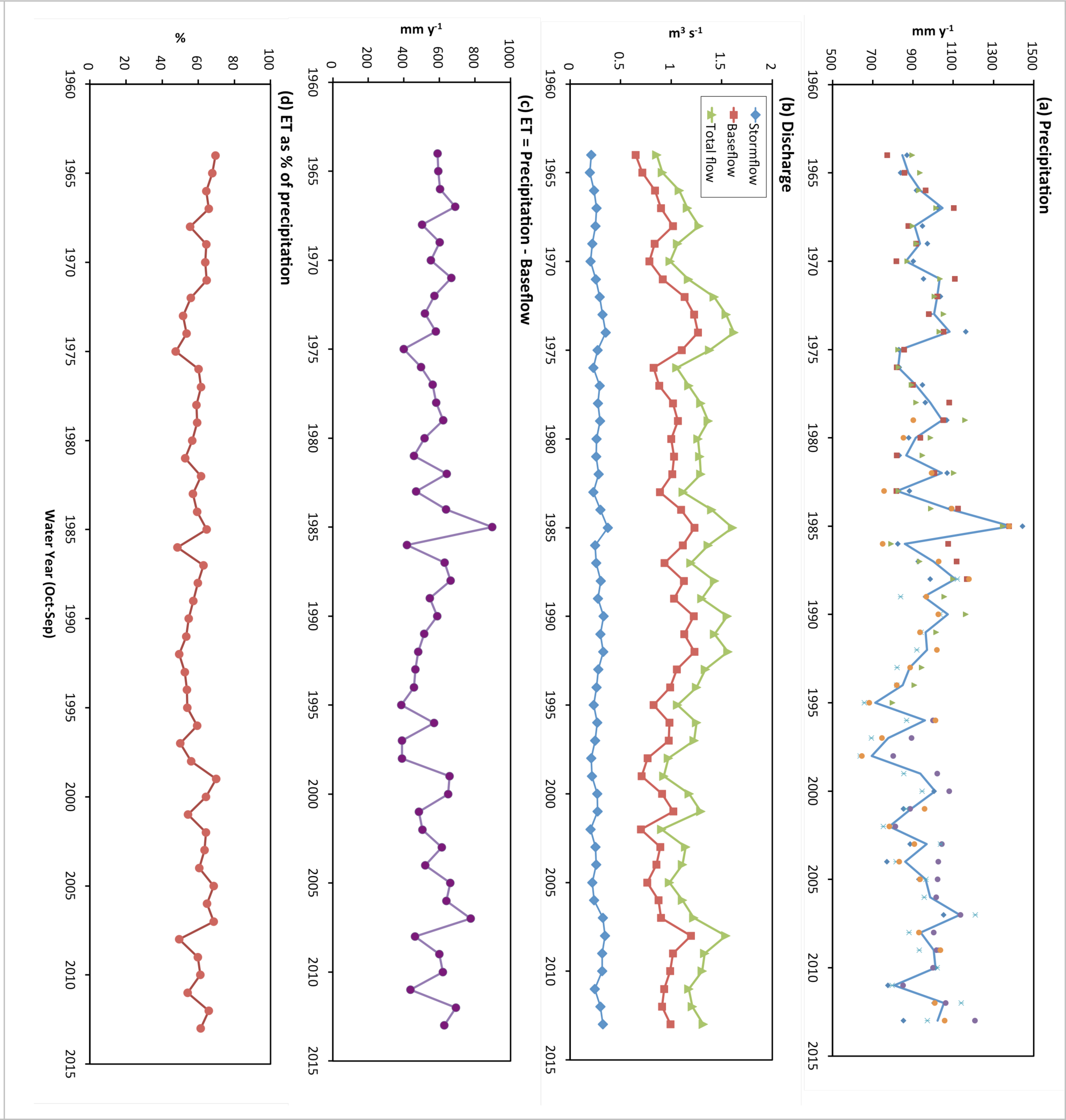
Precipitation, stream discharge, and evapotranspiration (ET). Panels show annual (Oct-Sep) values of (a) precipitation measured at 3-6 stations (mean = blue line); (b) stream discharge partitioned into baseflow and stormflow; (c) evapotranspiration (ET) estimated as the difference between precipitation and baseflow discharge; and (d) ET as a percentage of annual precipitation.

One or more of the four meteorological variables that control atmospheric evaporative demand—wind speed, atmospheric humidity, net radiation, and air temperature—could have changed over the 50 y, as global- and continental-scale analyses have indicated significant changes in these variables in recent decades (Wild, 2009; Willett *et al.,* 2008; McVicar *et al.,* 2012b). The effects of changes in these variables on atmospheric evaporative demand could be to enhance or counteract each other, and the resultant effect on ET is particularly important where evaporation is limited by energy rather than water (e.g., decreasing wind speeds tend to counteract the effect of increasing temperatures: McVicar *et al.,* 2012a, b). The region has experienced a 1.14°C increase in mean annual air temperature (50-y mean = 8.95°C) which in turn equates to a 0.90 millibar (mb) increase in saturated vapor pressure (50-y mean = 13.5 mb) over the 50-y period (Figure 4). Consistent data across the study period for wind speed, atmospheric humidity, and net radiation are not available for this locale.

**Figure 4.**
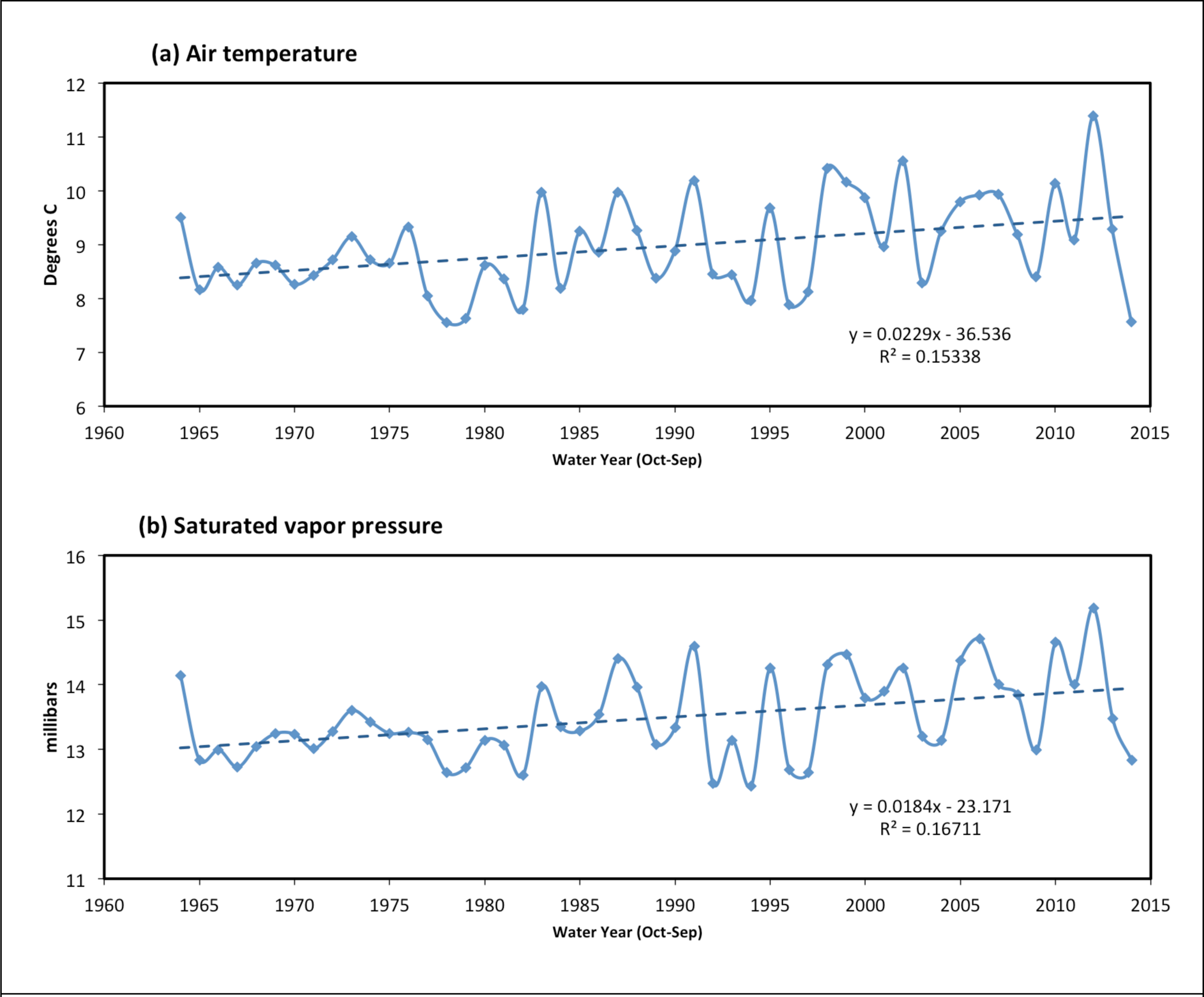
Air temperature (a) and saturated vapor pressure (b) for the Augusta Creek watershed, derived from the Midwest Regional Climate Center database (http://mrcc.isws.illinois.edu/). The positive changes are significant (P = 0.005 and 0.003 for temperature and vapor pressure, respectively) and amount to 1.14°;C and 0.90 millibars over the 50 years.

### 3.2 Catchment hydrology

Stream discharge partitioned into stormflow and baseflow shows how groundwater dominates the total flow of Augusta Creek; baseflow averaged 78% of the total discharge (Figure 3b). There is no linear trend in total (p = 0.14), stormflow (p = 0.91), or baseflow (p = 0.83) discharge over the 50 y. In this catchment, stormflow likely reflects mainly precipitation falling on lakes and wetlands that are contiguous with the stream channels because upland soils are highly permeable and there are few impervious surfaces and little overland runoff from uplands to the streams. This is supported by the comparison of annual stormflow volumes to the annual precipitation falling on contiguous lakes and wetlands: on average, stormflow amounts to 57% (range, 44-73%) of the precipitation with no linear trend over the 50 y (p = 0.09, data not shown). The balance, which equates to a mean of 408 mm y^-1^, could largely be explained by evapotranspirative losses from the lakes and wetlands. If stormflow originating as overland flow from the uplands were important, the total stormflow volume would exceed the precipitation on lakes and wetlands.

Our annual water balances for Augusta Creek resemble earlier estimates calculated by Rheaume (1990) over three representative years (1971, 1977 and 1985), which indicated that 62, 65 and 59%, respectively, of the annual precipitation was returned to the atmosphere as ET, mainly during the growing season (May–Sep), although those estimates included ET from contiguous lakes and wetlands as well as uplands. That study also employed hydrograph separation to estimate that about 75% of the annual stream flow in those years was supported by groundwater discharge; our estimate of mean baseflow contribution over the 50-y period is 78%.

Our estimate of ET, based on the difference between precipitation on the upland catchment and baseflow discharge out of the catchment, averaged 563 ± 103 mm y^-1^ (Figure 3c), with no linear trend (p = 0.98). Expressed as a percentage of annual precipitation, ET averaged 59 ± 6% (Figure 3d), also with no trend over the 50 y (p = 0.88). Therefore these data show that ET from upland areas of the Augusta Creek catchment has remained remarkably stable over the past 50 y in spite of large changes in land cover towards less area in annual crops and more in deciduous forest.

### 3.3 ET rates from representative vegetation types

We estimated ET in annual crops and perennial vegetation over the 2009–2014 period from high-resolution changes in soil water profiles (Figure 5). Except for 2012, which was a drought year, mean growing season ET rates were 495 ± 48 mm y^-1^ for maize, 524 ± 79 for grasslands (fallow and prairie), and 532 ± 47 for woody vegetation (deciduous forest, shrubland, and poplar). These rates are statistically indistinguishable among vegetation types (p>0.05), further supporting the hypothesis that ET rates are similar among annual crops, perennial grasslands, and forests in the Augusta Creek catchment. These ET observations span years of varying warmth (Figure 4) but show no relationship with mean growing-season temperature.

**Figure 5.**
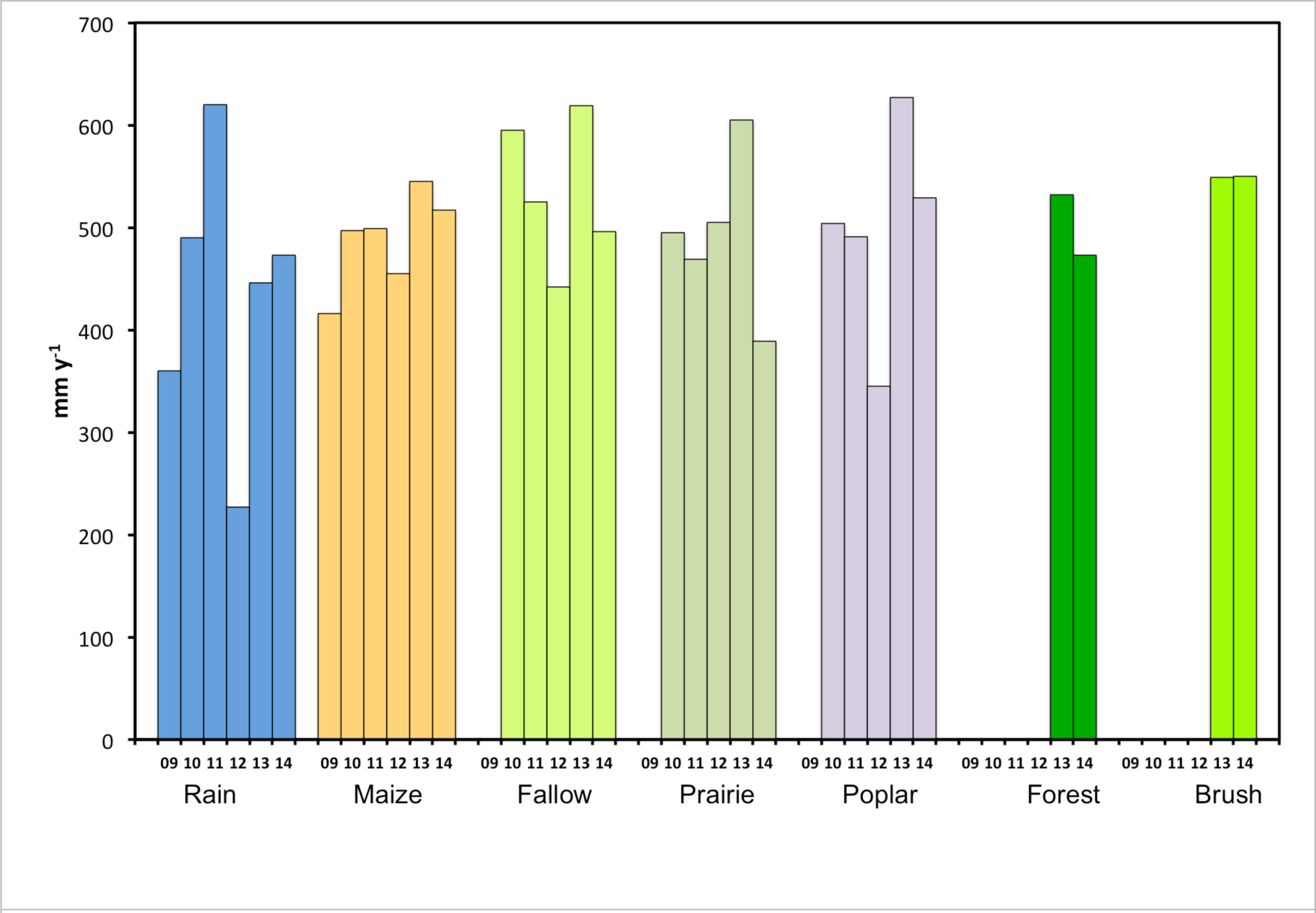
Rainfall (blue) and evapotranspiration over the growing season (2009-14) from annual maize and herbaceous and perennial vegetation, estimated from continuous observations of plant water uptake in soil profiles.

While soil water-based ET rates, excluding the 2012 drought year, are lower than the catchment-based ET rates of 600 ± 59 mm y^-1^ in those years (2009, 2010, 2011, 2013, and 2014 in Figure 3c), the soil water-based ET estimates reflect only the growing seasons. Year-round eddy covariance measurements of water fluxes in maize and grasslands at KBS indicate that about 30% of ET occurs outside the May-Sep growing season (Abraha *et al.,* 2015). Adding 30% to the soil water-based ET rates brings rates for maize, grasslands, and woody vegetation to 643, 681, and 692 mm y^-1^, respectively, all higher but within 15% of the catchment-based ET measurements over those years.

## 4. Discussion

### 4.1 Possible explanations for the stability of ET

There are several possible explanations for the long-term stability of catchment ET that we believe are unlikely. One is that there may not have been sufficient time for hydrologic responses to be detected. While the mean transit time for groundwater movement in this kind of catchment is likely greater than a decade (e.g., Saad, 2008), groundwater discharge rates from an unconfined and connected aquifer system would respond to changing recharge at far faster time scales (McDonnell and Beven, 2014). Succession from grassland to forest can be protracted, but the MIRIS forest cover data indicate that most of the reforestation occurred in the first 14 years of the study period (i.e., 1964–78). Many long-term paired catchment studies have shown that water yield after regrowth of harvested forest tends to approach a stable rate within about 10–25 years (Hornbeck *et al.,* 1993; Brown *et al.,* 2013).

Another possibility is that the degree of land cover change over the study period (27% of the upland catchment abandoned from annual crops and 20% of it becoming reforested; Figure 2) may not be sufficiently large to signal a change in water yield, even if annual crops and perennial vegetation had large differences in ET rates. Again, this is unlikely because long-term paired catchment studies have shown significant change with as little as 20% of the catchment either deforested or afforested (Brown *et al.,* 2005).

Also possible is that there are offsetting effects exerted by different land covers in the vicinity (Albertson *et al.,* 2001; van Dijk *et al.*, 2012), but this does not seem likely because adjacent catchments have similar mosaics of land cover, and the entire region has experienced similar changes in vegetation over this time period. Compensating land use changes that result in no net change in ET are also a possibility, such as the changes in crops grown as noted above. However the ET rate of oats that were commonly grown in the 1960s and 1970s is unlikely to differ much from the maize that replaced them (Allen *et al.,* 1998).

Over the past 50 y the mean annual air temperature has increased by about 1.14 °C (Figure 4), and the frost-free season has become longer by about 9 days (Kunkle, 2015). Evapotranspiration could increase with warming if available water were not limiting, other meteorological changes did not offset the temperature effect (McVicar *et al.*, 2012b), and the vegetation could remain active over the longer growing season. However, during the growing season when most (∼70%) of the ET occurs, available soil water typically becomes limiting to ET (Hamilton *et al.,* 2015). Also, most annual crops and many grasses would senesce before the end of the potential growing season because their development is regulated by degree-days (Parmesan and Hanley, 2015).

Extrapolation of observations from small catchments that are entirely covered by one kind of vegetation to complex mixtures of vegetation may not be as straightforward as it would seem. Models of ET and discharge from catchments with mixed land covers has often proven challenging to validate, and a variety of possible reasons have been considered (van Dijk *et al.,* 2012). Methodological issues identified by those authors include uncertainties in land cover, precipitation and discharge data; in the case of the Augusta Creek catchment, however, the precipitation and land cover data are likely to be quite accurate. It is also possible that other catchment climate characteristics that we have not considered are more influential to ET than land cover (e.g., Wilcox and Huang, 2010). Physical explanations noted by van Dijk *et al.,* (2012) for poor model performance include recirculation of intercepted rainfall, which tends to be more important in forests, and lateral water redistribution between vegetation types; identifying the potential importance of these physical explanations in the Augusta Creek catchment is beyond the scope of this study.

We cannot rule out the possibility that changes in the meteorological drivers of atmospheric water demand (i.e., temperature, net radiation, wind speed, and atmospheric humidity; McVicar *et al.*, 2012a, b) could have offset the effects of land cover changes on ET. The steadily increasing partial pressure of atmospheric carbon dioxide could also have reduced plant transpiration rates, although its effect on ET is most pronounced in warm, highly water-limited (i.e., arid and subarid) regions (Donohue *et al.*, 2013; Yang *et al.*, 2016; Trancoso *et al.*, 2017). In any case the offset of land cover effects on ET by these atmospheric changes would be a regional phenomenon contributing to the resilience of catchment ET and discharge.

### 4.2 Conclusion

Evapotranspirative water loss in the upland portion of the Augusta Creek catchment has been remarkably resilient across a 50-y period of decreasing cropland, increasing perennial vegetation cover, and warming temperatures, leaving a relatively consistent proportion of precipitation for groundwater recharge and streamflow. Our ET estimates based on catchment water balances compare well with direct measurements in the same catchment since 2009 based on soil water monitoring by time-domain reflectometry for grasslands, annual crops, and perennial bioenergy crops and forest. These observations suggest that water use by rainfed annual crops and perennial vegetation is similar in this setting, and that in humid catchments with soil permeability little affected by land cover, catchment water balances are not likely to be very sensitive to near-term future changes in land cover and climate as long as the land is vegetated, and crops are not irrigated. One such land cover change could be an increase in the cultivation of perennial herbaceous crops for biofuel production, which, based on our findings, does not seem likely to alter catchment water balances in this kind of setting.

## Supplementary Information

**Figure S1.**
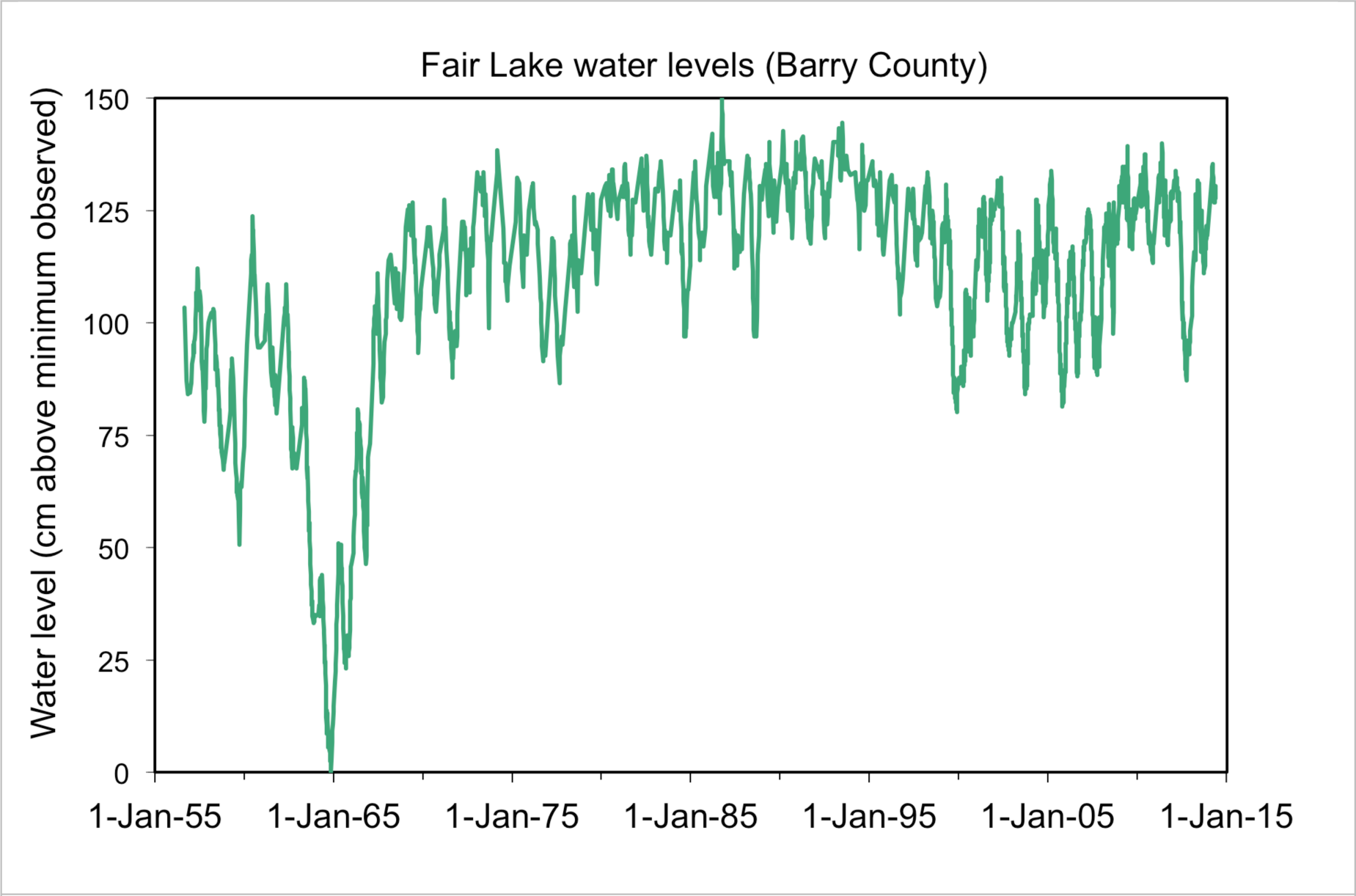
Water levels in Fair Lake, one of two lakes forming the headwaters of the Augusta Creek system. Note that these data extend back to well before the start of the study period on 1 Oct 1964, and encompass a series of drought years in the early 1960s. Since 1967 there has not been a unidirectional trend across years that would suggest large changes in groundwater or surface water storage. No other local lakes, whether draining to streams or isolated, are known to have changed unidirectionally over the study period. Data are available at http://lter.kbs.msu.edu/datatables/381.

**Figure S2.**
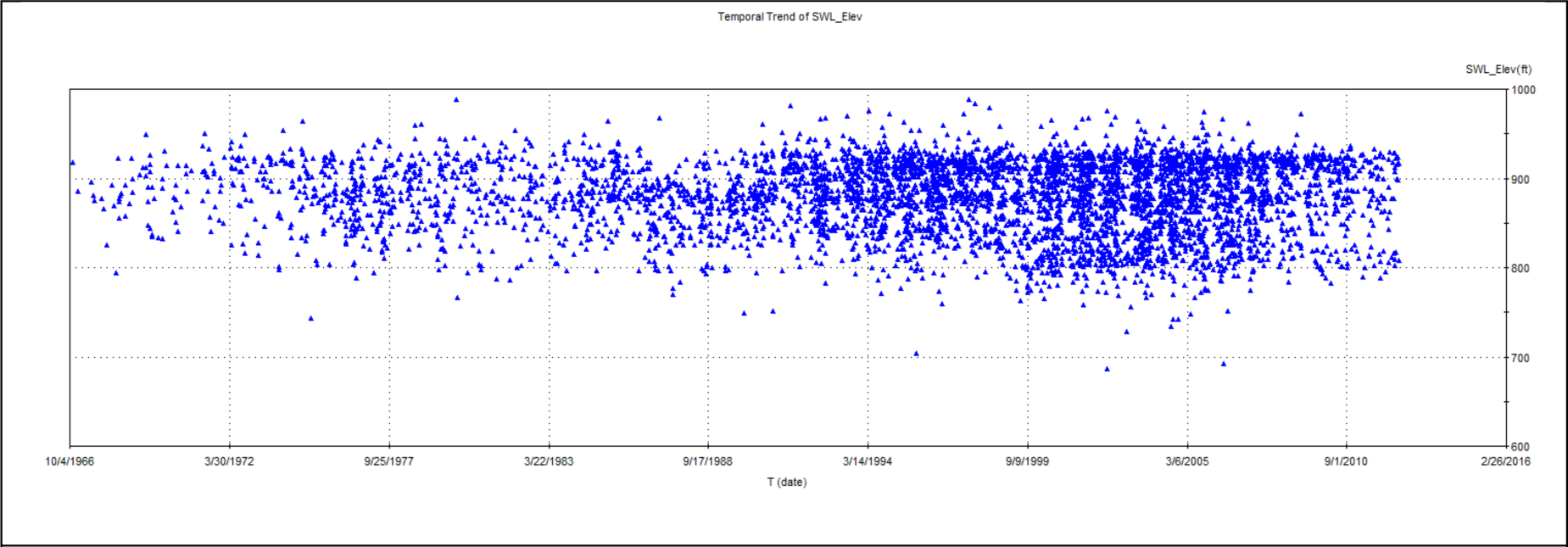
Static water levels measured upon installation of residential water supply wells in the vicinity of the Augusta Creek watershed. Data compiled from public records by Shu-Guang Li of Michigan State University.

**Figure S3.**
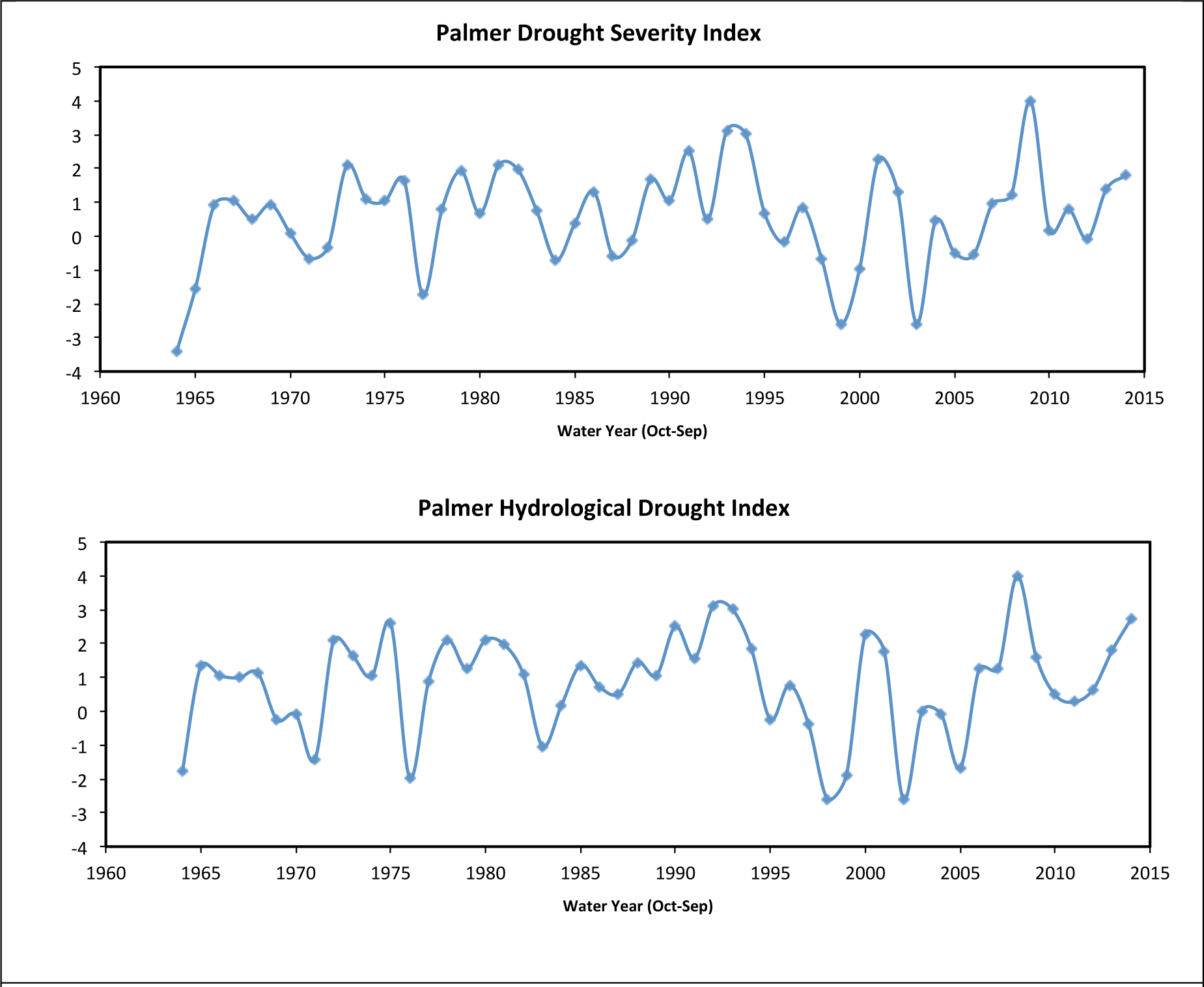
The Palmer Drought Severity Index (a) and the Palmer Hydrological Drought Index (b) for the region encompassing the Augusta Creek watershed, derived from the Midwest Regional Climate Center database (http://mrcc.isws.illinois.edu/). There is no significant linear trend in either index (p = 0.34 and 067, respectively).

## Acknowledgments

We thank A.K. Bhardwaj, S. Bohm, K. Kahmark, and S.-G. Li for instrumentation and data assistance, local citizens W. Shafer, T. Smith, and W. Knollenberg for precipitation data supplemental to that from our research and National Weather Service stations, and the numerous people at Michigan State University and the U.S. Geological Survey who helped maintain the precipitation and stream discharge records since 1964. T. McVicar, J.J. McDonnell and T. Dunne read earlier versions and provided helpful advice on data interpretation. Financial support for this work was provided by the U.S. Department of Energy through the Great Lakes Bioenergy Research Center (DOE BER Office of Science DE-FC02-07ER64494 and DOE OBP Office of Energy Efficiency and Renewable Energy DE-AC05-76RL01830), the U.S. National Science Foundation (LTER program, DEB 1027253), and the Michigan Agricultural Experiment Station.

